# Molecular packing structure of fibrin fibers resolved by X-ray scattering and molecular modeling

**DOI:** 10.1101/2020.01.15.907253

**Authors:** Karin A. Jansen, Artem Zhmurov, Bart E. Vos, Giuseppe Portale, D. Hermida Merino, Rustem I. Litvinov, Valerie Tutwiler, Nicholas A. Kurniawan, Wim Bras, John W. Weisel, Valeri Barsegov, Gijsje H. Koenderink

**Affiliations:** AMOLF, Biological Soft Matter group, Amsterdam, The Netherlands; UMC Utrecht, Department of Pathology, 3508 GA Utrecht, The Netherlands; KTH Royal Institute of Technology, Stockholm, Sweden; Sechenov University, Moscow 119991, Russian Federation; Institute of Cell Biology, Center of Molecular Biology of Inflammation, University of Münster, Münster, Germany; Macromolecular Chemistry and New Polymeric Materials, Zernike Institute for Advanced Materials, University of Groningen, Nijenborgh 4, 9747 AG Groningen, The Netherlands; Netherlands Organization for Scientific Research (NWO), DUBBLE CRG at the ESRF, 71 Avenue des Martyrs, 38000 Grenoble Cedex, France; Department of Cell and Developmental Biology, Perelman School of Medicine, University of Pennsylvania, Philadelphia, Pennsylvania, USA; Institute of Fundamental Medicine and Biology, Kazan Federal University, 18 Kremlyovskaya St., Kazan 420008, Russian Federation; Department of Biomedical Engineering and Institute for Complex Molecular Systems, Eindhoven University of Technology, Eindhoven, The Netherlands; Chemical Sciences Division, Oak Ridge National Laboratory, One Bethel Valley Road, Oak Ridge Tennessee, 37831, United States; Department of Chemistry, University of Massachusetts, 1 University Ave., Lowell, MA, 01854, USA; Department of Bionanoscience, Kavli Institute of Nanoscience Delft, Delft University of Technology, Van der Maasweg 9, 2629 HZ Delft, the Netherlands

**Keywords:** filamentous proteins, biopolymers, self-assembly, hemostasis, full-atom simulations

## Abstract

Fibrin is the major extracellular component of blood clots and a proteinaceous hydrogel used as a versatile biomaterial. Fibrin forms branched networks of polymeric fibers, built of laterally associated double-stranded protofibrils. This multiscale hierarchical structure is crucial for the extraordinary mechanical resilience of blood clots. Yet, the structural basis of clot mechanical properties remains largely unclear due, in part, to the unresolved molecular packing structure of fibrin fibers. Here we quantitatively assess the packing structure of fibrin fibers by combining Small Angle X-ray Scattering (SAXS) measurements of fibrin networks reconstituted under a wide range of conditions with computational molecular modeling of fibrin oligomers. The number, positions, and intensities of the Bragg peaks observed in the SAXS experiments were reproduced computationally based on the all-atom molecular structure of reconstructed fibrin protofibrils. Specifically, the model correctly predicts the intensities of the reflections of the 22.5 nm axial repeat, corresponding to the half-staggered longitudinal arrangement of fibrin molecules. In addition, the SAXS measurements showed that protofibrils within fibrin fibers have a partially ordered lateral arrangement with a characteristic transverse repeat distance of 13 nm, irrespective of the fiber thickness. These findings provide fundamental insights into the molecular structure of fibrin clots that underlies their biological and physical properties.

---

Fibrin forms the polymeric mechanical scaffold of blood clots and thrombi. Fibrin networks, along with platelets and erythrocytes, serve to seal sites of vascular injury and promote wound healing^1^. In addition, fibrin hydrogels have been extensively used as biomaterials in tissue engineering due to their unique biological and physical characteristics, such as porosity, deformability, elasticity and biodegradability^2^. Fibrin networks have a complex hierarchical structure that is crucial for their material properties^3^. At the network scale, fibrin fibers form a branched, space-filling elastic network that is able to withstand large mechanical deformations exerted by flowing blood, platelet-induced contraction, and deformations of the vessel wall. The fibers are in turn made up of laterally associated protofibrils^4^, which themselves are two-stranded linear filaments of half-staggered fibrin monomers^5^. The molecular packing of fibrin fibers, together with the structural flexibility of individual fibrin molecules, results in remarkable mechanical properties. For example, an individual fibrin fiber can be stretched by about twice its length before changes in elastic properties occur^6-7^. Whole fibrin clots are similarly deformable and resilient^8-10^. Furthermore, the molecular packing of fibrin fibers affects the susceptibility of fibrin to enzymatic lysis, a process called fibrinolysis that dissolves fully or partially obstructive blood clots (thrombi) to restore impaired blood flow^11-12^.

The structure of fibrin networks has been studied at various structural levels. The structure of the monomers has been elucidated by X-ray crystallography^13^, electron microscopy^14^ and atomic force microscopy^15-16^ and has been modeled by full-atom molecular dynamics (MD) simulations^17-21^. The structure and assembly mechanism of fibrin protofibrils has likewise been studied by electron microscopy and atomic force microscopy^22-24^ and through full-atom MD simulations^22, 25^. The structure at the network level has been characterized mostly by light scattering^23, 26-28^, confocal light microscopy^29-30^, and scanning electron microscopy^31^.

The least well understood aspect of the hierarchical structure of fibrin is the three-dimensional molecular packing of protofibrils within fibrin fibers. Along the fiber axis, fibrin monomers are known to be half-staggered, as evident from cross-striations with a 22.5-nm repeat in high-resolution images of fibers taken by electron or atomic force microscopy^14-16, 23^. Transverse to the fiber axis, it is less clear how protofibrils are organized. Most of what we know about the 3D packing structure of fibrin fibers comes from small angle X-ray scattering (SAXS) because it allows the molecular arrangement of fibrin fibers to be probed in their native hydrated state. Early on, SAXS was used to demonstrate the half-staggered molecular packing structure of fibrin fibers, which shows up in the form of a Bragg peak at a wave vector corresponding to the 22.5 nm repeat distance^14, 32^. Later measurements by SAXS^4, 32-35^ and neutron diffraction^36^ confirmed that the axial packing order is long-ranged, and reasonably well-ordered, along a fiber, as evident from the presence of quite narrow peaks corresponding to higher-order reflections of the 22.5-nm axial repeat distance. Surprisingly, while the third and higher-order reflections are often observed in the SAXS spectra, the second order reflection is usually missing or very faint^32, 36-37^, although the origin of this peak suppression is unclear. The suppression of such a low angle peak usually indicates a higher symmetry or a form factor scattering that has a minimum at this *q* value, but so far the source of this long range ordering has remained elusive.

Transverse to the fiber axis, the packing of protofibrils inside the fibrin fiber was shown by SAXS to be less ordered than the axial packing, but there is controversy about the degree of disorder. Some X-ray scattering studies concluded that there was no order in the lateral packing of fibrin fibers^4, 32^, while others found evidence of considerable lateral crystallinity^34, 36, 38^. It was proposed that fibrin fibers might be partially crystalline as a consequence of defects in the radial packing structure, and that the degree of order might depend on the conditions of fibrin self-assembly^39^. The lateral packing order might also vary for fibers with different dimensions^14^.

One of the main challenges in resolving the three-dimensional packing of protofibrils within fibrin fibers in SAXS experiments is the lack of experimental data about the detailed structure of the protofibril, because it is a short-lived intermediate oligomer formed at the early stages of fibrin formation. This makes it difficult to relate the positions, intensities, and widths of low angle diffraction peaks, which for brevity we will refer to as ‘Bragg peaks’, observed in SAXS spectra to structural signatures originating from molecular packing within fibrin fibers. As a powerful methodological advantage, here we provide a structural model-based interpretation of experimental SAXS spectra for fibrin based on the full-atomic modeling of fibrin protofibrils *in silico*. By coupling experiments and modeling, we resolve the axial and radial packing arrangements of fibrin fibers. The theoretically reconstructed SAXS spectra captured both the number, positions and intensities of the Bragg peaks observed by experimental SAXS and explain the long-standing riddle as to why the second order reflection of the 22.5 nm axial repeat is suppressed. By systematically varying the fibrin fiber thickness, we found that the range of axial order reflected by the width of the Bragg peaks in the SAXS spectra of isotropic fibrin networks is determined by the protein packing density of the fibers. Furthermore, our results confirm predictions of proposed model that the radial packing of fibrin fibers is partially ordered^39^ with a newly determined characteristic repeat distance of 13 nm that is independent of the fiber thickness.

## RESULTS

### Probing the axial packing of fibrin fibers with SAXS

To probe the sensitivity of the small angle X-ray diffraction to subtle differences in the fiber structure, we performed SAXS measurements on isotropic fibrin networks assembled using different polymerization conditions inside glass capillaries (Fig. 1A). By changing the fibrinogen concentration and buffer composition (see Materials and Methods), we varied the fiber thickness, expressed as the average number of protofibrils per cross-section of a fibrin fiber (*N*_*p*_), over a wide range from *N*_*p*_ = 2 up to *N*_*p*_ = 435, as determined by light scattering measurements (see Table 1 and confocal reflectance images in Fig. 2B-D). Light scattering measurements showed that the variation in fiber thickness was accompanied by strong variations in the protein mass density of the fibers, as summarized in Table 1. The mass density ranged from *ρ* = 11.6 mg/ml for *N*_*p*_ = 47 up to *ρ* = 248 mg/ml for *N*_*p*_ = 368. This observation is qualitatively consistent with earlier reports of strong variations in fibrin fiber density with assembly conditions^39-41^. Note that for the *N*_p_ = 2 networks we could only determine the mass-length ratio (and thus *N*_p_) of the fibers from light scattering and not the fiber radius (and thus *ρ*), because these thin fibers have radii of only 7.5-15 nm, smaller than the wavelength of light^9^.

**Figure 1.**
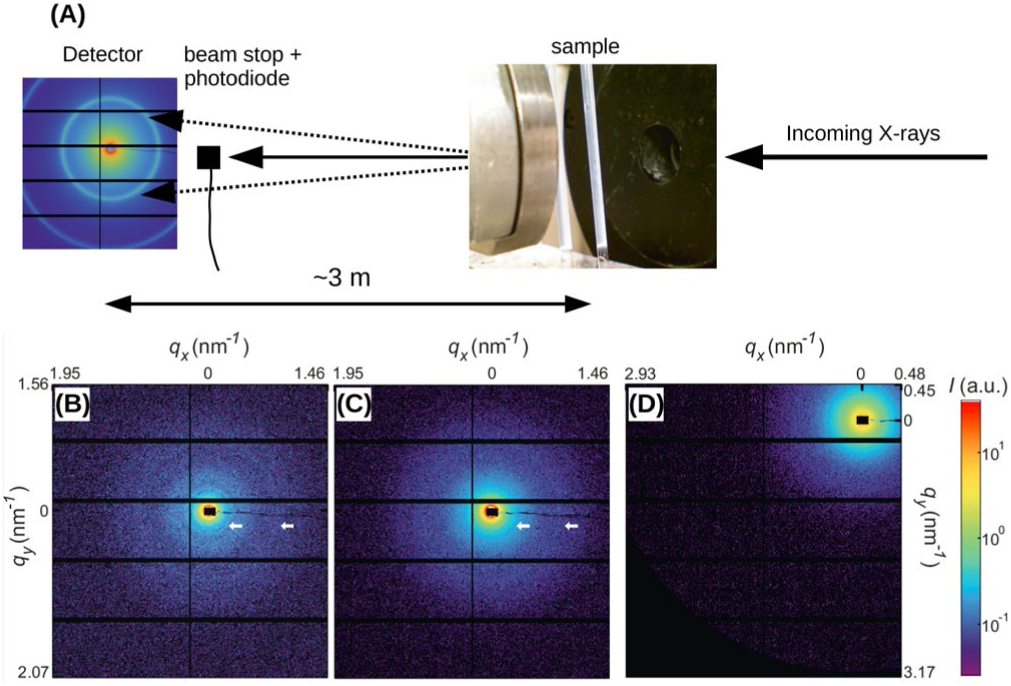
(A) Schematic depiction of the SAXS experiment. Fibrin samples in glass capillaries (photo on the right) are illuminated with an X-ray beam. The small-angle diffraction pattern (left, example pattern recorded for a powdered silver behenate crystal) is captured with a camera placed a distance of 3 meters away from the sample. A beam stop blocks the unscattered X-ray beam and a photodiode records the intensity. (B) Two-dimensional SAXS pattern after background subtraction for a fibrin network composed of thick and dense fibers (*N*_*p*_ = 368, *ρ* = 248 mg/ml), (C) for thick fibers of somewhat lower mass density (*N*_*p*_ = 435, *ρ* = 190 mg/ml), and (D) for thin fibers (*N*_*p*_ = 2, *ρ* ≈ 3.4 mg/ml). White arrows in (B) and (C) indicate the first and third order reflections of the axial packing periodicity of fibrin (22.5 nm); in (D) these peaks are not distinguishable from the background. Note that the detector was translated in the *x-* and *y-*directions in between the recordings of (B,C) and (D). The black rectangle in the center of the SAXS patterns represents the beam stop. Intensity is in arbitrary units (see the color bar).

**Figure 2.**
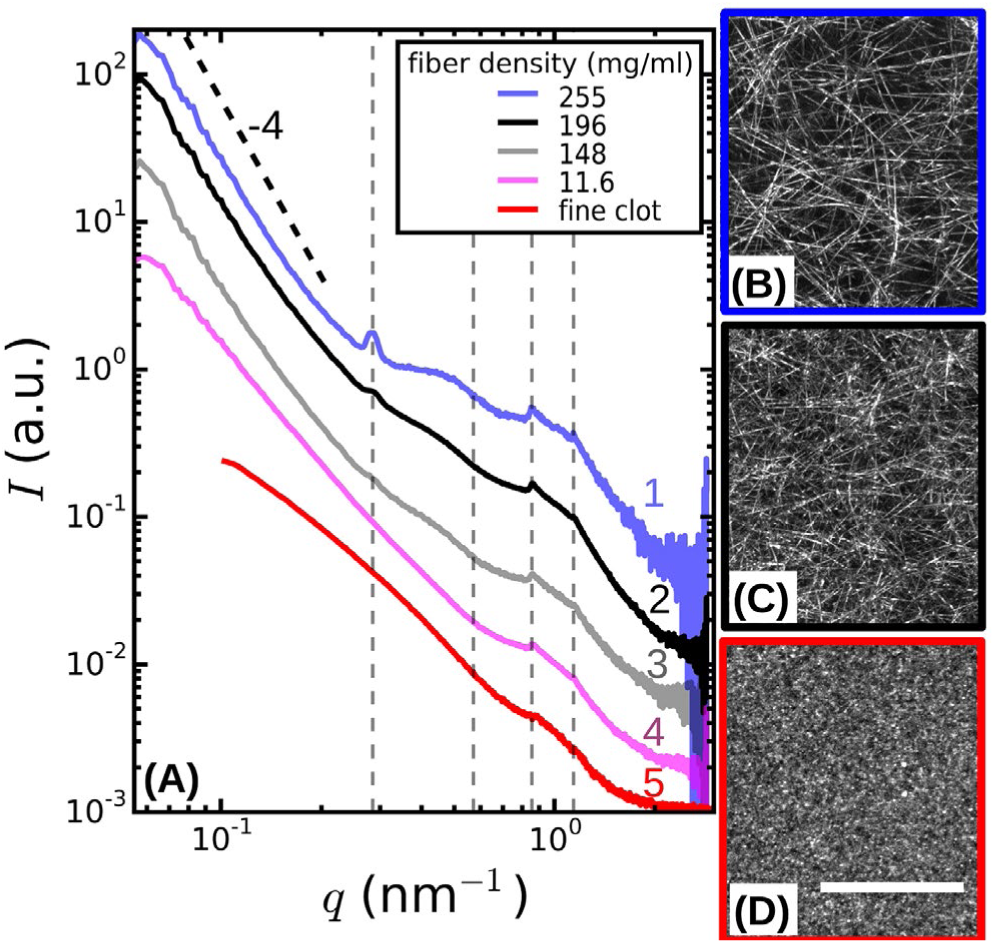
(A) SAXS spectra obtained by radial integration of the 2D SAXS patterns of fibrin networks with fibers of varying protofibril number N_p_. For fine clots (red curve), *N*_p_ =2. The other curves represent coarse (thick-fiber) clots with different protein mass densities *ρ* as specified in the legend (see Supporting Information Table S1). The curves are shifted along the *y*-axis for clarity. Vertical dotted lines indicate the expected positions of the first, second, third and fourth order reflections of the axial packing distance (22.5 nm), while the dotted black line indicates the Porod scattering regime where *I*(*q*) ∝ *q*^-4^ for reference. (B-D) Confocal reflectance images of coarse fibrin networks with fiber mass densities of 248 and 144 mg/ml, and for fine clots. Images show maximum intensity projections over a total depth of 20 μm. The scale bar is 20 μm.

The 2D SAXS patterns of fibrin were isotropic in all cases, showing concentric reflection rings (cf. white arrows in Figure 1B-C), as would be expected for a random fibrous network. The intensities of the reflections were larger for thick fibers (*N*_*p*_ = 368 and 435 in Fig. 1B and 1C, respectively) than for thin fibers (*N*_*p*_ = 2 in Fig. 1D). The *N*_p_-dependence of the scattering was more evident from the SAXS 1D profiles obtained by the radial integration of the 2D scattering patterns (Figure 2A). For the fibers with the highest mass density (*ρ* = 248 mg/ml) we observe a clear Bragg peak at a wave number *q* = 0.285 nm^-1^ (labeled as *curve 1*). This peak corresponds to a repeat distance of 22.2 nm and it can thus be assigned as the first-order reflection of the fibrin half-staggered axial packing repeat^32^. In case of long-range axial packing order, higher-order reflections are expected at *q* = 0.57 (second), *q* = 0.855 (third) and *q* = 1.14 nm^-1^ (fourth) (denoted by dashed vertical lines). Interestingly, we only observe the third and fourth order reflections. For fibers with a slightly lower mass density (*ρ* = 191 mg/ml), we again observed first, third and fourth order reflections, but with a smaller peak height (*curve 2* in Figure 2A). The same reflections are still faintly visible for even less dense fibers (*ρ* = 144 mg/ml, *curve 3*). For the thinnest fibers with *ρ* = 35 mg/ml (*curve 4*) or *ρ* = 3.5 mg/ml (*curve 5*), the first order reflection is not visible at all, but we can still faintly discern the third and fourth order reflections. Qualitatively, these observations are consistent with prior SAXS measurements, where the first order reflection was clearly observed for fibrin networks composed of thick fibers^8, 34, 36, 38-39, 42-43^, but not for networks of protofibrils formed at high ionic strength^43^. The absence of the second order reflection from the SAXS spectra is also consistent with earlier SAXS studies^32, 39^, but the reason for the suppression of this peak is not clear.

### Molecular origin of fibrin SAXS spectra

To understand the molecular origin of the Bragg peaks in the SAXS spectra of fibrin, we performed a structure-based theoretical reconstruction of SAXS spectra using a complete atomic model of the two-stranded fibrin protofibril, a fibrin oligomer with 10 fibrin monomers in one strand and 9 in the other complementary strand (abbreviated as FP10-9). First, we computed the theoretical SAXS spectra using the distribution of atomic pair distances *p*(*r*) for protofibril fragment FP10-9, which is directly accessible from the all-atom MD simulations (Figure 3, *curve 2*). Here, the dashed vertical lines indicate the expected positions of the first, second, third and fourth order reflections of the half-staggered repeat distance. Consistent with the experimental data, the second order reflection was absent in the theoretical spectrum for fragment FP10-9. Hence, the complete structural model of two-stranded fibrin protofibrils FP10-9 accounts for the suppression of the second order reflection observed in the SAXS experiments.

**Figure 3.**
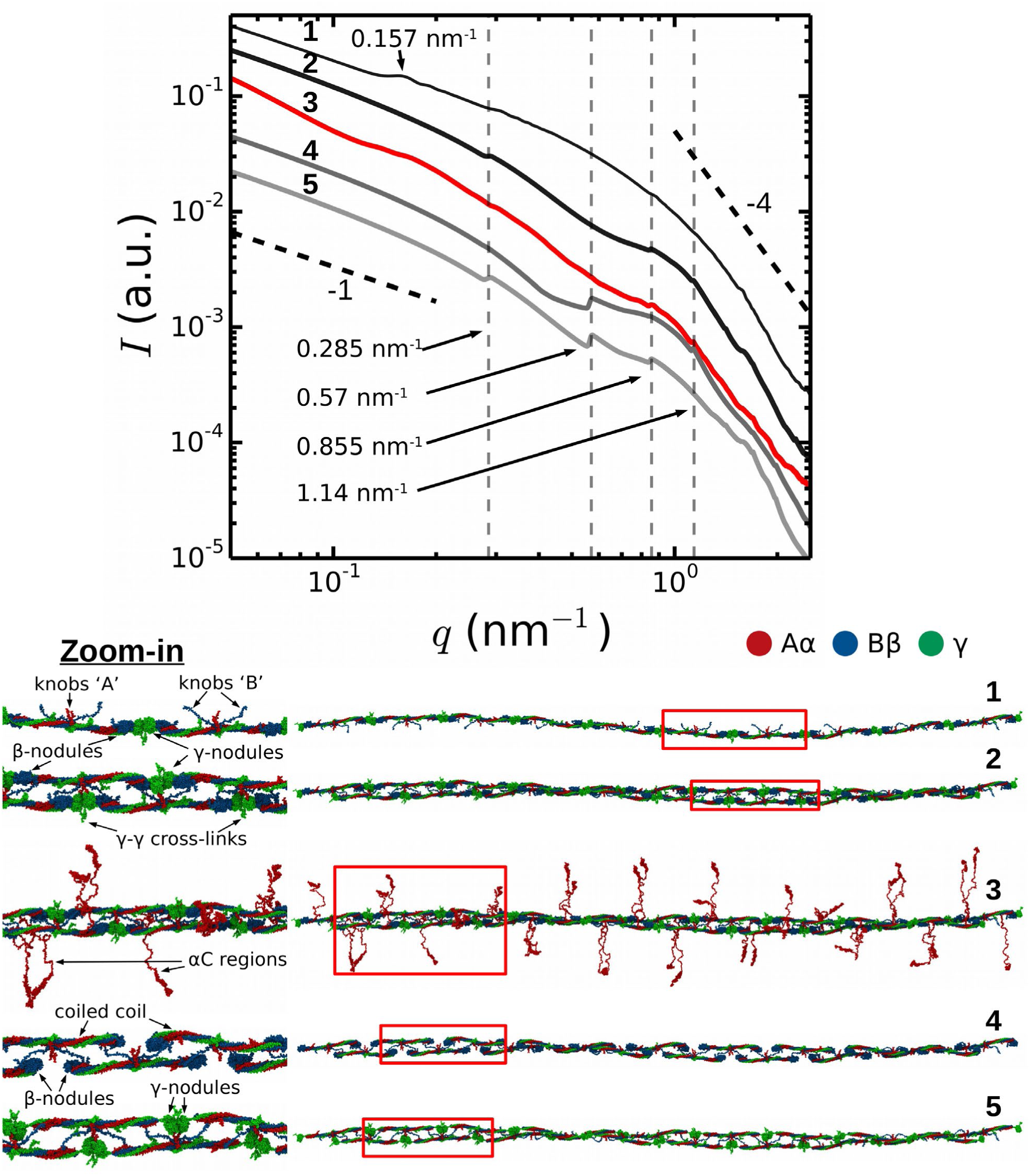
Theoretically reconstructed SAXS spectra (top) based on full-atom structural models of a fibrin protofibril and its variants (bottom). Numbers correspond to: (1) single-stranded fibrin polymer Fn_m_ with *m* = 10 fibrin monomers connected longitudinally at the D:D interface; (2) twisted double-stranded protofibril without the αC regions; (3) same protofibril as in 2, but with αC regions (shown in red) incorporated in a random coil conformation; (4) same structure as in 2, but with the γ-nodules removed; (5) same structure as in 2, but with the β-nodules removed. The black dashed lines indicate a power law with exponent −1 for low *q*-numbers (expected for cylinders, see Fig. S2 in the Supporting Information) and −4 for high *q*-numbers (corresponding to the Porod regime). Red boxes in the structural models in the bottom indicate regions where we zoomed in (displayed on the left) to show more structural detail.

Next, to understand the origin of the suppression of the second order reflection, we systematically modified the protofibril structure by removing selected important structural elements, i.e. the αC regions (FP10-9/αC), γ-nodules (FP10-9/γ), and β-nodules (FP10-9/β), and recalculating the corresponding SAXS spectrum for each truncated construct. The results presented in Figure 3 clearly demonstrate that the second order reflection is suppressed altogether in all SAXS spectra for fragment FP10-9 containing αC regions (*curve 3*) and without αC regions (*curve 2*). Nevertheless, the presence of αC regions does influence other reflections corresponding to the half-staggering distance, causing a reduced peak intensity and increased peak width. By contrast, when either all the γ-nodules (FP10-9/γ, *curve 4*) or β-nodules (FP10-9/β, *curve 5*) are removed from the protofibril structure, the second order reflection appears.

As an additional control, we also calculated a SAXS spectrum for a hypothetical single-stranded fibrin polymer, composed of a linear strand of 10 monomers connected end-to-end through the D:D self-association interface (Figure 3; *curve 1*). In this case, a new peak appeared at *q* = 0.157 nm^-1^ corresponding to a typical distance of 40 nm between the centers of mass of the two symmetrical lateral globular D regions. Each fibrin monomer is ∼45-nm long and has two globular ends of 6 nm each^13, 44^. Therefore, the distance between the centers of mass of the two globular ends of the molecule is 40 nm (see Supporting Information Fig. S3). This 0.157 nm^-1^ peak is suppressed for the double-stranded fibrin protofibrils. Hence, we conclude that the second order reflection is not present in the SAXS spectra of fibrin due to destructive interference from the symmetric nature of fibrin monomers. This resolves a long standing puzzle in the interpretation of X-ray scattering data from fibrin fibers.

### Probing the radial packing of fibrin fibers with SAXS

In addition to the Bragg peaks discussed above, the *q*-dependence of the X-ray scattering patterns contains information on the radial (transverse) packing structure of fibrin fibers. However, this information is more difficult to infer from the spectra than the axial packing order because the radial packing is more disordered. To determine the *q*-range where the SAXS spectra are sensitive to the radial packing structure, we first consider the length scales covered by the SAXS spectra based on an analytical model proposed by Ferri and co-workers^26-27, 29^ that treats the network as a collection of “blobs” of size *ξ*_*blob*_ with network fractal dimension *D*_*m*_ (see schematic in Fig. 4). Since fibrin fibers are rigid on scales much larger than the mesh size, we assume that each blob contains fiber segments that can be approximated by cylinders of a certain length (*l*) and diameter (*d*). In the absence of any internal packing structure, this model predicts three distinct scattering regimes as a function of *q* (black curve in Figure 4). At small *q*, the spectrum should exhibit a peak at *q* = *q*_1_, which is related to the long-range network order^45^. The position of this peak is inversely proportional to the average blob size^26-27^ according to *q*_1_ ≃ 4.4/*ξ*_*blob*_. In the middle range, *q*_1_ < *q* < *q*_2_, the scattering intensity is determined by the fractal network structure within the blobs, i.e. *I* ∝ *q*^-*Dm*^. Finally, in the limit of large *q* > *q*_2_, the Porod regime is reached. In this regime, scattering occurs from the interface between the fibers and the solvent. Hence, the onset of the Porod regime is set by the fiber diameter^26-27, 39^ according to *q*_2_≃2.2/*d*. In case of smooth filaments that lack internal structure, the scattering intensity is expected to decay as a power law with *q* with an exponent *α*_*s*_ = −4.

**Figure 4.**
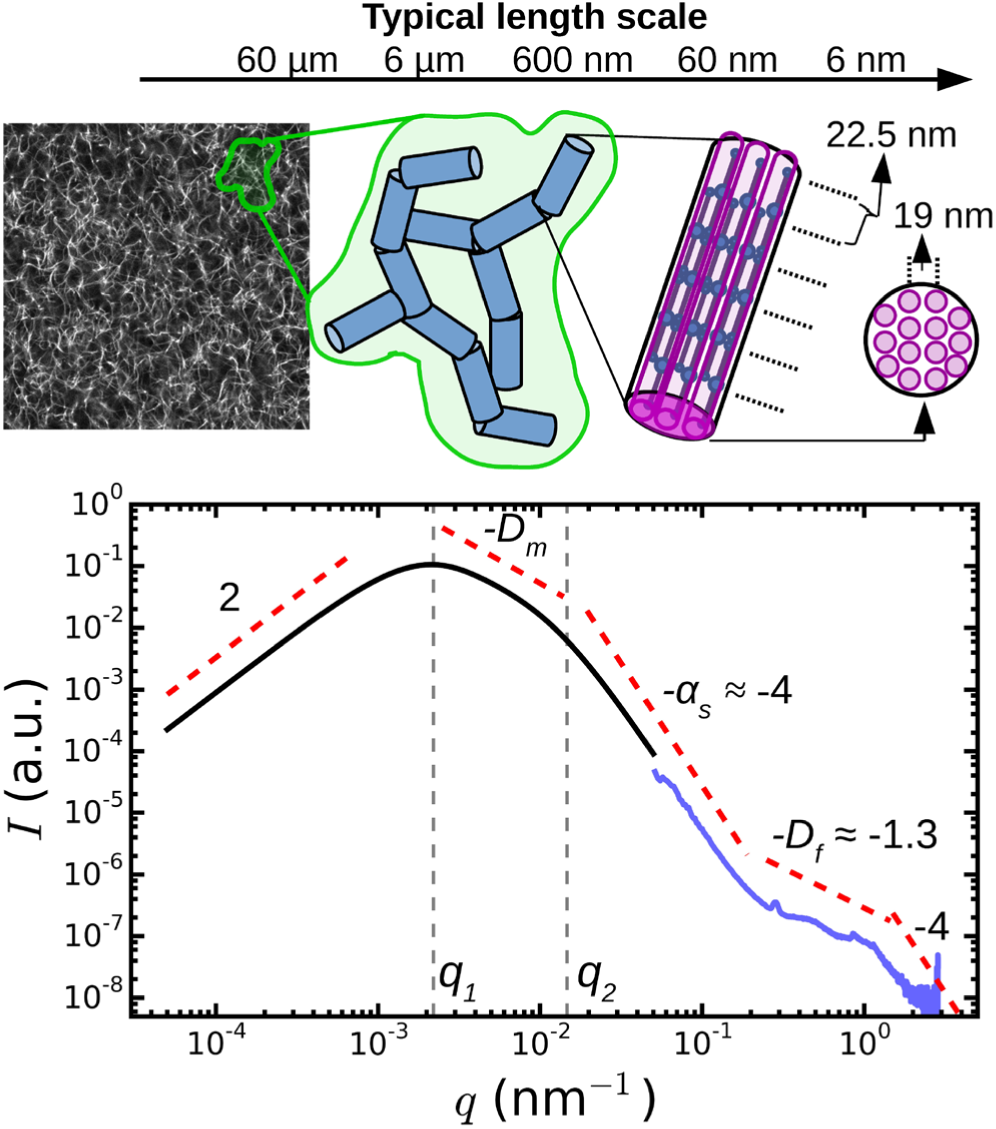
Model to distinguish between network and fiber structural features in light/X-ray scattering data. On large length scales (*q* < 10^−1^ nm^−1^), fibrin networks behave as a collection of blobs (shown in green in the schematic) of rigid fiber segments (blue). For fibers lacking any internal structure, this model predicts three *q*-regimes (black curve). Experimentally, SAXS curves (blue curve, for *N*_*p*_ = 368/*ρ* = 248 mg/ml) show additional fine structure at high *q*, which therefore provides information on the internal structure of the fibers. The model curve was calculated from the model of Ferri and co-workers^27, 29^, using as input a blob size *ξ*_*blob*_ = 10 μm, fiber segment length *l* = 0.5 μm, fiber diameter *d* = 0.2 μm, fractal dimension *D*_*m*_ = 1.3, and Porod exponent *α*_*s*_ = 4. The two curves are shifted along the *y*-axis for clarity (note arbitrary units). The top-left image shows a typical confocal fluorescence image of a fibrin network (100 μm × 100 μm).

When we compare this calculation with SAXS spectra measured for thick fibrin fibers (*N*_*p*_ = 368, blue curve in Fig. 4), we see that the low *q*-regime (*q* < 0.2 nm^-1^) of the SAXS spectrum corresponds to the Porod regime, where the scattering intensity decreases as *I* ∼ *q*^-4^. However, the high *q*-regime (*q* > 0.2 nm^-1^) of the spectra displays features not predicted by the scattering model. It is this regime that carries information about the internal structure of the fibers, and that we will consider below. Note that for networks of very thin fibers, with *N*_*p*_ = 2 (*curve 5* in Figure 2), we do not observe the Porod regime. Indeed for these thin fibers, the Porod regime should start only once *q* reaches ∼1 nm^-1^. In this case, the SAXS measurements can therefore not access the internal structure of the fibers, but they only access the *q*-regime where scattering is determined by the fractal structure of the network (*I* ∼ *q*^*-Dm*^). We find a value for the network fractal dimension *D*_m_ of around 1.5.

To infer the radial packing structure of the fibers from the high-*q* regime of the SAXS spectra, we need to assume a structural model for the cross-sectional packing. So far three different models have been proposed (Figure 5A). The first model assumes that protofibrils are packed in an ordered lattice^46^ that should give rise to a Bragg peak at a *q*-vector corresponding to the average spacing between the protofibrils. Indeed, some measurements by SAXS^35, 38^, SANS^36^, neutron diffraction^42, 47^ and energy-dispersive X-ray diffraction^34, 38^ revealed broad Bragg peaks, indicating partial disorder, but other SAXS studies found no evidence for lateral crystalline order^32^. The second model instead assumes that the fibers are completely disordered assemblies of protofibrils (Figure 5A), which should give rise to a power-law decay of the scattering intensity *I*∝*q*^*-Df*^ with *D*_f_ the fractal dimension of the fibers characterizing their radial mass distribution^48^. Physically, this means that fibrin fibers would have a dense core and a loose periphery with sparsely arranged protofibrils. This model is supported by measurements of the fluorescence intensity^49^ and stiffness^40^ of fibrin fibers as a function of their diameter. The third model is essentially a hybrid of the other two models, because it considers the lateral packing as crystalline but with defects (Figure 5A)^39^. This arrangement would result in a superposition of a fractal-like scattering pattern with broad peak(s) due to locally crystalline regions.

**Figure 5.**
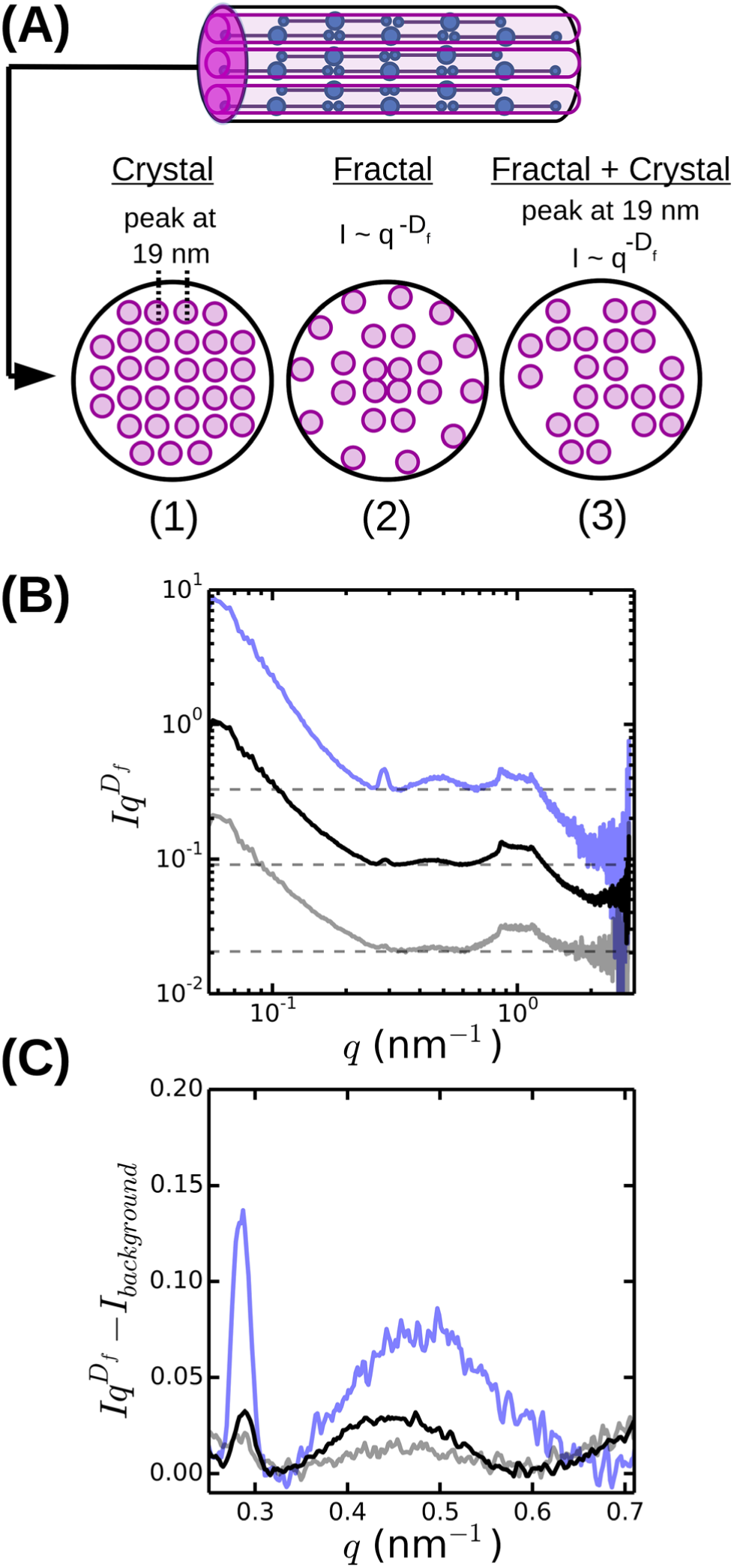
Radial (cross-sectional) packing structure of fibrin fibers. (A) Current models assume either (1) a crystalline array^46^ with a unit cell measuring 19×19×46 nm, or (2) a disordered structure^49^ characterized by a fractal dimension *D*_*f*_ = 1.3 characterizing the internal mass distribution, or (3) a superposition of a crystalline and a fractal structure, with disorder stemming from missing protofibrils^39^. (B) Rescaled SAXS spectra for fibrin networks with varying bundle size, obtained by multiplying SAXS spectra with *q*^*Df*^, reveal that fibrin fibers combine a fractal structure with partial radial order (broad peak at q ≈ 0.47 nm^-1^). (C) Magnified views of a small region of the rescaled spectra after subtraction of the background level (horizontal dashed lines in B). Spectra are shown for networks with *ρ* = 248 mg/ml (blue), 191 mg/ml (black) and 144 mg/ml. Spectra in (A) and (B) were shifted along the y-axis for clarity.

The spectrum shown in Fig. 4 is most consistent with the third model: we observe a superposition of an overall *I* ∼*q*^*-Df*^ power-law decay with an exponent *D*_*f*_ close to −1.3 (indicated by the red dashed line between 0.2 and 1 nm^-1^) with a broad and weak superimposed bump centered around *q* = 0.47 nm^-1^. This is more visible when we multiply the scattering intensities by *q*^*Df*^ and adjust *D*_*f*_ to make the resulting curves flat between *q* = 0.2 and 1 nm^-1^ (see Figure 5B). The best-fit values for *D*_*f*_ vary between 1.3 and 1.7 (Figure 6A), consistent with prior measurements based on SAXS^39^, fluorescence microscopy^49^, and single-fiber stretching^40, 50^. As shown in Figure 5C, the Iq^*D*f^-curves nicely reveal the narrow Bragg peak at *q* = 0.29 nm^-1^ corresponding to the axial half-staggered packing order of fibrin, as well as a much broader peak centered around *q* = 0.47 nm^-1^. We observe the broader peak only for the thick fibers (*N*_*p*_ = 291, 368, and 435) and not for the thinner fibers (*N*_*p*_ = 2 and 47). In view of the full-atom molecular modeling described above, it is unlikely that this peak is the second order peak originating from the axial packing. We therefore propose that it corresponds to lateral packing order, with a characteristic repeat distance between protofibrils of 13 nm. This spacing is in reasonable agreement with previous studies reporting repeat distances of ∼19 nm based on scattering methods or electron microscopy^14, 34-38, 42^. As shown in Figure 6B, we find that both the peak for axial order (solid circles) and for lateral order (open squares) are independent of the fiber thickness and the associated mass density.

**Figure 6.**
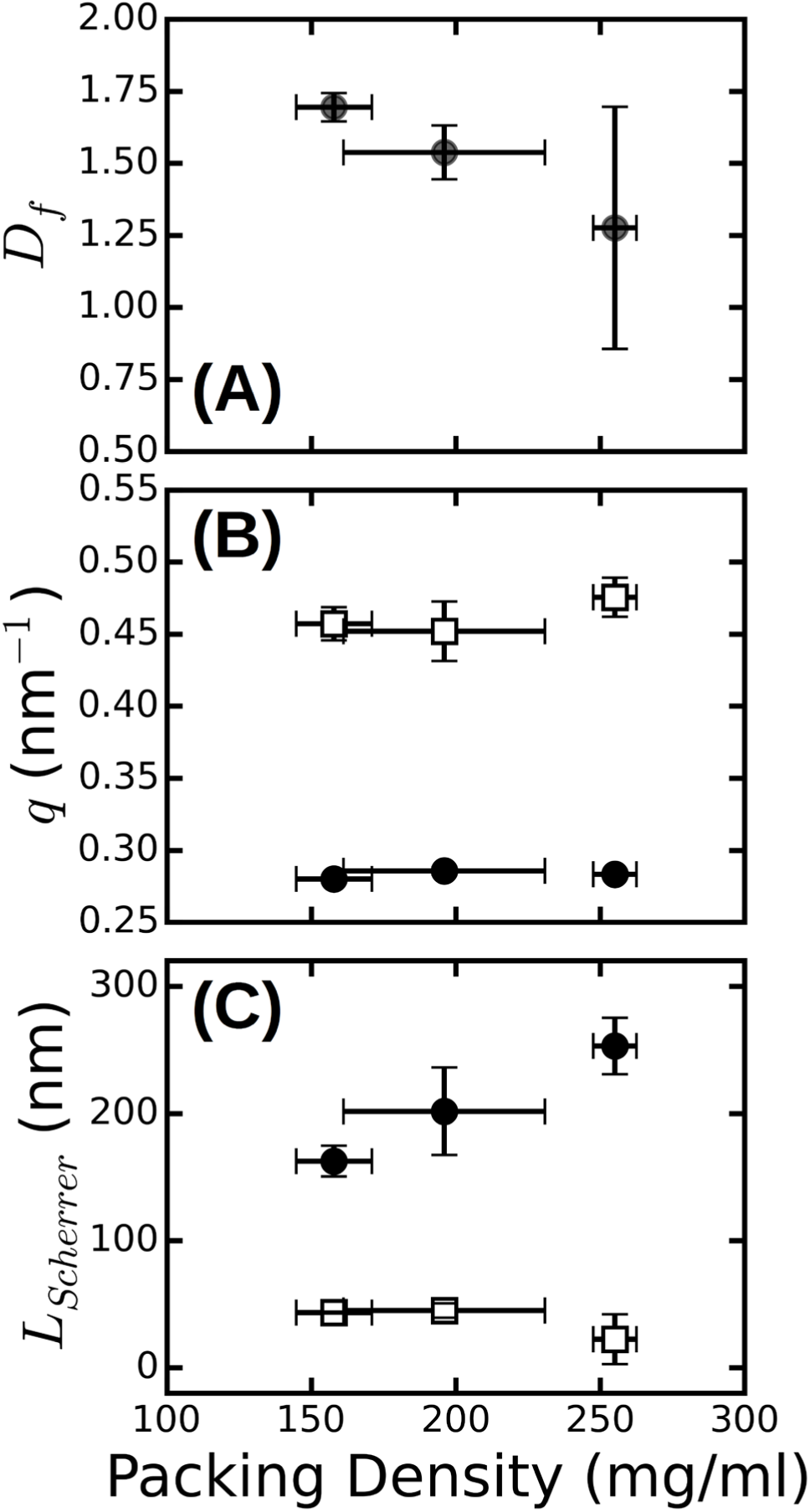
Radial and axial packing order of fibrin fibers. (A) Fiber fractal dimension *D*_*f*_ obtained from SAXS spectra in the range of 0.2 ≲*q* ≲1 nm^-1^, plotted as a function of the protein mass density *ρ*, determined by turbidimetry. (B) The *q*-positions of the first order Bragg peak corresponding to the axial packing periodicity of fibrin fibers (black solid circles), and of the broad peak, which we attribute to the radial packing order (open squares). The peak positions are independent of fiber mass density. (D) Corresponding Scherrer length, computed from the full width at half maximum of the peaks. The Scherrer length characteristic of axial order (black solid circles) shows a stronger dependence on fiber mass density than the Scherrer length characteristic of radial order (open squares). Error bars represent the standard deviation from fitting data recorded at multiple positions over at least two repeat measurements.

The large width of the *q* = 0.47 nm^-1^ peak suggests that the radial ordering is only short-ranged. Indeed, the fiber diameter poses a strict upper limit on the range of order. In a perfectly crystalline system, the average crystal size can be approximately quantified by the Scherrer peak broadening, which sets the full width at half maximum of a Bragg’s peak Δ*q*, i.e. *L*_*scherrer*_ = 2*π*/Δ*q*^51^. We used this expression to estimate the apparent crystal size as a function of fiber protein density from the width of the peak determined by Gaussian peak fits. As shown in Figure 6C (open squares), *L*_*scherrer*_ was ∼40 nm and independent of fiber density. This observation suggests that the extent of (crystalline) cross-sectional ordering is limited not by the fiber diameter, but rather by the intrinsic disorder. The presence of intrinsic disorder is consistent with the existence of a fractal dimension, and also agrees with reported electron microscopy data^14^. By contrast, the Scherrer length associated with the axial packing periodicity systematically increases with increasing fiber mass density (Figure 6D, solid circles). This finding suggests that the axial ordering becomes more pronounced as the fibrin fibers become more densely packed.

## DISCUSSION

Here, we have investigated the internal packing structures of fibrin fibers by performing SAXS measurements on fibrin networks with structural variations induced by changing the fibrin assembly conditions. To interpret the results in terms of the molecular packing structure, we complemented the experimental SAXS measurements with SAXS patterns derived *in silico* based on full-atom reconstructions of fibrin protofibrils.

We have shown that the SAXS spectra of fibrin networks reveal both longitudinal and lateral (i.e. axial and radial) molecular packing within fibrin fibers. The axial packing gave rise to a sharp reflection at *q*=0.285 nm^-1^ and several higher-order reflections (Bragg peaks) indicative of long-range 22.5-nm packing periodicity of fibrin. This periodicity corresponds with findings in a number of X-ray and neutron scattering experiments^35-36, 38, 42, 47^ as well as electron microscopy^14, 23^ and atomic force microscopy^15-16^ of individual fibrin fibers. Our work advances this well-known structural characteristic of fibrin assembly and structure by revealing that the axial packing in fibrin fibers is sensitive to the fiber protein density. In particular, these axial reflections are only seen when the fibers are thick enough, i.e., the number of protofibrils per cross-section, *N*_*p*_, is larger than ∼40. Moreover, the denser is the fiber, the more intense and narrow the first order Bragg peak. Strikingly, the second order reflection of the axial packing order is missing in all fibrin samples tested. With the help of full-atom modeling of fibrin protofibrils (Figure 3), we were able to determine that the origin of this phenomenon is the symmetry arising as a result of the spacing of the γ- and β-nodules. The second order reflection shows up only after either all the γ- or all the β-nodules are computationally removed from the fibrin molecules that form protofibrils. Accordingly, the appearance of the second order reflection could serve as a signature of potential forced unfolding transitions in fibrin networks under tensile mechanical stress, given that simulations proposed primary unfolding of the γ-nodules^17-18^. Furthermore, the SAXS simulations show that the disordered αC regions emanating from the protofibrils broaden the Bragg peaks.

We observed a systematic increase of the apparent axial packing order with increasing mass density of the fibers. A possible explanation is that the densest fibrin fibers in our study were prepared from gel-filtered fibrinogen (see Methods) containing non-aggregated monomeric fibrinogen molecules of uniform molecular length. In support of this explanation, the presence of fibrinogen aggregates in the clotting mixture has been shown to alter the kinetics of polymerization, impair the assembly of monomers into protofibrils and fibers^52^ and reduce the protein density of fiber fibers and lessen coherency of the 22.5-nm axial packing^53^. Another possible source of variability in the fibrin fiber packing and their structural arrangement is that the presence of the fibrinogen γ’ splicing variant in plasma and plasma-drived fibrinogen preparations affects the number of protofibrils and protein density within fibers^54^.

Whether protofibrils within a fibrin fiber are ordered or disordered in the direction transverse to the fiber axis has been a matter of debate. Our results show that fibrin fibers are partially ordered with elements of a fractal structure, giving rise to a power-law decay of the scattering intensity with *q*, and of a crystalline-like structure, with a broad superimposed peak centered around *q* = 0.47 nm^-1^. The peak position reveals a 13-nm repeat distance between protofibrils, close to values reported elsewhere^14, 34, 36-38, 42^. We found that the fractal dimension of the fibers is on the order of *D*_f_ = 1.5, well within the range of other reported fractal dimensions for fibrin obtained by atomic force microscopy and SAXS^39-40, 49^. However, we find that the fractal dimension is dependent on the fiber mass density. To the best of our knowledge, no one has looked at the fractal dimension of the fibrin fiber with systematic changes in the packing densities experimentally. However, an increase of fractal dimension with increasing protein density can be expected based on the model presented by Yeromonahos et al^39^. In this model, the lateral packing of a fibrin fiber is effectively fractal because not all the crystalline positions are used by protofibrils (Figure 5A). In the context of this model, an increase in fiber protein density means physically that more empty spaces in the crystal structure are filled by protofibrils, bringing the overall structure closer to a homogeneous packing with a limiting *D*f value of 2.

## CONCLUSIONS

To get insight in the structural hierarchy of fibrin clots, we have performed SAXS measurements on fibrin networks composed of fibers with varying thickness and internal protein mass density in combination with full-atom simulations of protofibrils. We investigated the effects of axial and lateral packing on the scattering patterns by varying the fiber protein density from 3.4 mg/ml up to 250 mg/ml. We show that the Bragg peaks corresponding to axial order are much more pronounced for denser fibers and that the peak width also depends on the fiber protein density. We show, for the first time, that the axially symmetric molecular packing structure of fibrin can explain the suppression of the second order reflection of the 22-nm axial repeat observed here as well as in previous publications. When we artificially perturb the symmetry by computational removal of the γ- or β-nodules, the second order peak appears. An interesting prediction of this model is that SAXS could be used to probe whether the γ-nodules unfold upon fibrin network deformation, since unfolding should result in the appearance of the second order peak. Finally, we show that fibrin fibers have a partially ordered lateral packing with a characteristic repeat distance of 13 nm, independent of the fiber thickness or the number of protofibrils per fiber cross-section. Our findings demonstrate that SAXS in combination with computational modeling provides a powerful method to extract structural information at different spatial scales. In future, this approach may be used to study how fibrin networks respond to mechanical perturbations and to probe how mutations, polymorphisms, and posttranslational modifications of fibrinogen impact fibrin network structure.

## MATERIALS AND METHODS

### Formation of fibrin clots with varying structure

Human fibrinogen (FIB 3) and human α-thrombin were obtained from Enzyme Research Laboratories (Swansea, UK). FIB 3 was depleted from plasminogen, von Willebrand Factor, and fibronectin and delivered at a concentration of 13.64 mg/ml in a 20 mM sodium citrate-HCl buffer, pH 7.4 (stock solution). To vary the fiber thickness (quantified as the average number of protofibrils per cross-section of the fiber, *N*_*p*_) and fiber protein density *ρ* (in units of mass per volume), we used 4 different fibrin assembly conditions (see Table S1 in the Supporting Information):

1. *Fine clot conditions:* Networks with *N*_*p*_ = 2 and *ρ* ≈ 3.5 mg/ml were obtained from the fibrinogen stock preparation dialyzed against buffer with a high ionic strength and alkaline pH (50 mM Tris-HCl, 400 mM NaCl, pH 8.5). Fibrin was formed after diluting fibrinogen to the desired concentration in the same buffer supplemented with 3.2 mM CaCl_2_ followed by addition of thrombin to a final concentration of 0.5 U/mL and incubation for 2 hours at 37°C. These conditions were previously shown to result in so-called fine clots with extremely thin fibers due to a minimal degree of protofibril lateral association^55^.
2. *Assembly from as-received stock:* Networks with *N*_*p*_ = 47 and *ρ* = 34.9 mg/ml at a fibrinogen concentration of 8 mg/ml were obtained after diluting the fibrinogen stock preparation in the assembly buffer containing 20 mM HEPES (4-(2-hydroxyethyl)-1-piperazineethanesulfonic acid), 150 mM NaCl and 5 mM CaCl_2_, pH 7.4. Fibrin formation was initiated by adding 0.5 U/mL thrombin (final concentration) and the network was allowed to form for 4 hours at 37°C before measurements.
3. *Assembly from dialyzed stock fibrinogen preparation:* Fibrin networks with thicker fibers were obtained from the fibrinogen stock solution dialyzed against assembly buffer without CaCl_2_. We prepared networks with *N*_*p*_ = 291 and *ρ* = 144 by adding thrombin (0.5 U/mL final concentration) to fibrinogen at 4 mg/ml and networks with *N*_*p*_ = 435 and *ρ* =190 mg/ml by clotting fibrinogen at 8 mg/ml with the same amount of thrombin. The protein concentration after dialysis was determined using a NanoDrop spectrophotometer from the absorbance at a wavelength of 280 nm with correction for scattering at 320 nm as described^56^.
4. *Assembly from gel-filtered monomeric fibrinogen:* Networks with *N*_*p*_ =368 and *ρ* = 248 mg/ml at a 4 mg/ml fibrinogen concentration were prepared from monomeric fibrinogen obtained after removing contaminating fibrinogen oligomers from the fibrinogen stock solution by gel-filtration on Superdex-200 as described^52^.

### Confocal microscopy

Fibrin networks prepared as described above were imaged on a Nikon Eclipse Ti inverted confocal fluorescence microscope equipped with a 100× oil immersion objective lens (NA 1.49), a 488-nm laser (Coherent, Utrecht, The Netherlands) for illumination, and a photomultiplier tube detector (A1; Nikon, Amsterdam, the Netherlands). We collected stacks of confocal slices over a total depth of 10 μm and with a spacing of 0.125 μm. We performed confocal reflectance microscopy on unlabeled fibrin samples (see Fig. 2) and fluorescence imaging (see Fig. 4) on networks polymerized from unlabeled fibrinogen and Alexa488-labeled human fibrinogen (Life Technologies, Bleiswijk, the Netherlands) mixed in a 30:1 molar ratio.

### Turbidimetry

To characterize the fibrin fibers in terms of the number of protofibrils per fiber *N*_*p*_ and the protein mass density *ρ*, we performed turbidity measurements using a UV1 Spectrophotometer (Thermo Optek). Fibrin samples prepared as described in disposable plastic cuvettes (UV-Cuvette micro, Plastibrand) with an optical path length of 1 cm for low-turbidity samples and 2 mm for high-turbidity samples. The optical density (OD) for wavelengths between 400 and 900 nm with 1 nm intervals was measured relative to a reference sample consisting of just buffer. The turbidity (in units of cm^-1^) follows from the optical density by multiplication by ln(10) times the path length^39^. We analyzed the data using a custom-written Python script (available upon request) that fits the wavelength dependence of the turbidity to an analytical model describing light scattering of fibrous networks^28^ (see Fig. S1 in the Supporting Information for example measurements and fits). The model approximates the fibers as solid cylinders, but takes into account the fractal structure of branched fibrin networks as characterized by the network fractal dimension *D*_*m*_. The model also includes a correction for the wavelength dispersion of the solvent refractive index *n*_s_(*λ*) and the differential refractive index d*n*(*λ*)/d*c* (with *c* the protein mass concentration) based on Cauchy’s empirical relations^28^:

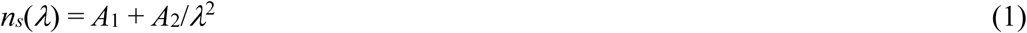

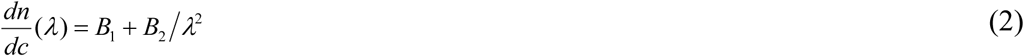

with optical constants *A*_1_ = 1.3270, *A*_2_ = 3.0595·10^−3^ µm^2^, *B*_1_ = 0.1856 cm^3^/g and *B*_2_ = 2.550·10^−3^ cm^3^µm^2^/g from Refs.^28, 57^. The model involves the two structural parameters characteristic of the fibers that we aim to determine, namely their radius *R* and mass-length ratio *ν*, and two parameters characterizing the network structure, namely its fractal dimension *D*_m_ and mesh size *ξ*. We separately determined *D*_m_ for each network by Fourier transforming the confocal reflection microscopy images, radially integrating the Fourier transformed images to calculate a power spectrum, and fitting the power spectrum to a power-law I(*q*) ∝ *q*^-*D*m^ over a range of spatial frequencies *q* corresponding to length scales between 0.6 μm (three times the diffraction limit) and 10 μm (1/10^th^ of the image size).^9-30, 60^ Resulting values for *D*_*m*_ varied between 1.4 and 1.5. The parameters *ξ* and *ν* are related according to:

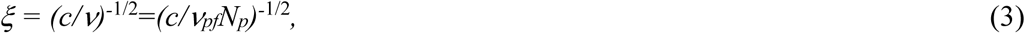

where *ν*_pf_ = 1.55·10^11^ Da/cm is the mass-per-length of a single double-stranded fibrin protofibril^58^. We therefore used an iterative process to find the best-fit values for *ξ, ν* and *R*. We first performed a fit using an approximate initial value for *ξ* as input that was based on a visual inspection of the confocal data. We then repeated the fitting procedure with *ξ* updated from the *ν*-value obtained from the preceding fit using Eq. (3) until *ξ* and *ν* changed by <1%, which typically required fewer than 5 iterations. We performed this fitting procedure for each sample separately, to account for possible sample-to-sample variations in *ξ*. From the fiber radius *R* and mass-per-length ν obtained from the fits, we calculated the fiber mass density *ρ =ν/πR*^***2***^. In case of fine clots, the fiber radius is smaller than the wavelength, so the wavelength-dependent turbidity was fitted to the Carr model for thin fibers^58^ to determine *ν*. The fiber radius was estimated in earlier work from our lab to be in the range of ∼7.5-15 nm based on electron microscopy imaging^9^.

### Small Angle X-ray Scattering of fibrin networks

Small Angle X-ray Scattering (SAXS) was performed at the DUBBLE Beamline (BM26B) of the European Synchrotron Radiation Facility (ESRF, Grenoble, France)^59^. The range of the wave vector *q* was calibrated using silver behenate powder as a standard. The sample-to-detector (P1M) distance was about 3 m and the energy of the beam was 12 keV. The beam dimensions on the sample were about 900×700 μm. The fibrin samples were prepared as described earlier inside 2-mm borosilicate glass capillaries with 0.01-mm wall thickness (Hilgenberg, Germany). Capillaries filled with buffer were used to determine background scattering. Since there can be variations in capillary thickness, we ensured that a background was taken for every capillary and on every spot we measured, before we polymerized the fibrin gels in the same capillaries.

### Atomic structural models of fibrin protofibrils

To interpret the experimental SAXS spectra, we used the complete atomic models of two-stranded fibrin oligomers with 10 fibrin monomers in one strand and 9 in the other complementary strand (abbreviated as FP10-9). The fibrin molecule is composed of three polypeptide chains denoted Aα, Bβ and γ, which fold into a symmetric trinodular structure that is 45 nm in length^60^. The central nodule is formed by the N-terminal portions of all the six chains, and is connected to the globular β- and γ-nodules formed by the C-terminal parts of the β- and γ-chains via triple α-helical coiled-coils^46^. Half-staggered assembly into protofibrils is driven by specific interactions between knobs ‘A’ in the central E region and holes ‘a’ in the lateral D-regions of adjacent fibrin molecules^5, 61^. The flexible C-terminal portions of the Aα chains known as the αC regions form compact αC-domains that are tethered to the molecule by flexible αC-connectors^62^. Fibrin oligomers were reconstructed *in silico* using CHARMM^63^, complete with A:a and B:a knob-hole bonds, γ-γ covalent crosslinking, and αC chains, as described elsewhere^25^. One of the general features to note is that in all structural models, the two strands inside the protofibril are twisted around one another, forming a superhelical structure. This is an important property that affects the lateral aggregation of protofirbrils into fibrin fibers and sets the limit of fibers’ thickness^64^. In this superhelical structure, fibrin monomers are slightly tilted with respect to the helical axis, which reduces the scattering periodicity.

To obtain the atomic model of a protofibril without the αC-domain (FP10-9/αC), the complete atomic structures were truncated at residue Gln221 in both α-chains in all 19 fibrin monomers. To reconstruct the virtual model of a protofibril without the γ-nodules (FP10-9/γ), both γ-chains in all fibrin monomers in both strands were truncated past residue Cys137. To obtain the structural model of a protofibril without the β-nodules (FP10-9/β), the β-chains were truncated in all 19 fibrin monomers starting from residue Cys197. Simulations were performed using the in-house codes MDis and SOP-GPU^18, 65^. The obtained *in silico* models of structural variants FP10-9/αC, FP10-9/γ, and FP10-9/β were energy-minimized using the steepest descent algorithm algorithm^66^. A single-stranded fibrin polymer Fn_m_ with *m* = 10 fibrin monomers connected longitudinally at the D:D interface was constructed using resolved crystallographic structures by connecting monomers end-to-end at the D-D junction. To build a single-stranded fibrinogen dimer, the double-D structure (PDB structure^67^ 1FZG) was aligned with the full-length fibrinogen (PDB structure 3GHG [13]) so that one of the D regions of the double-D fragment overlapped with one of the D regions of the fibrin molecule. We used the Kabsch algorithm^68^ to align the globular parts of the molecules. Here, we superimposed the Cα-atoms of resolved residues Bβ197-461 in the β-nodule and γ139-411 in the γ-nodule. The procedure was then repeated until the desired length of the oligomer was reached. Data visualization was done using VMD^69^.

### Theoretical reconstruction of SAXS spectra

The one-dimensional SAXS scattering spectrum *I(q)* of a collection of single fibrin fibers was calculated in the limit of a large number of scattering particles *N*, using the following relation:

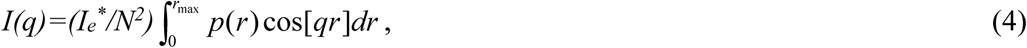

where *p*(*r*) is the distribution of atomic pair distances and *r*_max_ is the maximum particle-particle distance (i.e. fiber length). The distribution of atomic pair distances *p*(*r*) can be readily evaluated using the output from the MD simulations (structure files) by measuring the binary distances of atomic pairs. In Eq. (4), 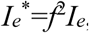, where *f* is the scattering strength and *I*_*e*_*= (I*_*0*_*e*^*2*^*/mc*^*2*^*) (1+*cos^*2*^*[2ψ])/(2d*^*2*^) is the intensity of a wave, scattered by a single electron^70-71^. Here, *I*_*0*_ is the intensity of an incoming wave, *d* is the distance from the object to the detector, *q=*|*q*|*=* 2*π*sin(*ψ*)/*λ* is the momentum transfer, λ is the wavelength of scattered radiation, and 2*ψ* is the scattering angle.

Since the fibrin networks are isotropic as evidenced from the isotropic SAXS patterns (Fig. 1B-D) and from the confocal images (Fig. 2B-D), we averaged the scattering intensity of single fibers over all orientations in 3D, according to:

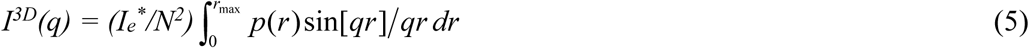

In the theoretical reconstruction of SAXS spectra for fibrin fibers, we calculated the distribution of atomic pair distances, *p(r)* and then computed *I*^*3D*^*(q)* by performing the numerical integration. The calculations were done using custom-written codes in C/C++/CUDA.

## ASSOCIATED CONTENT

## Supporting Information

A table summarizing the structural characterization of the different fibrin networks used in this study by turbidimetry; Figures showing examples of wavelength-dependent turbidity measurements on fibrin networks, a comparison of a full-atom simulation of FO6-5 protofibrils with the calculated form factor of a cylinder, and a schematic of a single stranded fibrin protofibril to explain the existence of twist inside a fibrin bundle.

## ACKNOWLEDGEMENTS

We thank Baldomero Alonso Latorre (AMOLF) for help with SAXS data analysis, and Federica Burla (AMOLF) and Fabio Ferri (Università dell’Insubria) for help with analysis of the turbidimetry data. This work was part of the research program of the Foundation for Fundamental Research on Matter (FOM), which is financially supported by the Netherlands Organization for Scientific Research (NWO). We gratefully acknowledge access to the DUBBLE BM26B beamline at the ESRF made possible by NWO. WB’s contribution is based upon work supported by Oak Ridge National Laboratory, managed by UT-Battelle, LLC, for the U.S. Department of Energy. This work was further supported by the American Heart Association grants 15GRNT23150000 and 13GRNT16960013, NIH grants HL135254 and UO1-HL116330, the NSF grants DMR 1505662 and DMR 1505316, and the Program for Competitive Growth at Kazan Federal University.

## TABLE OF CONTENTS GRAPHIC

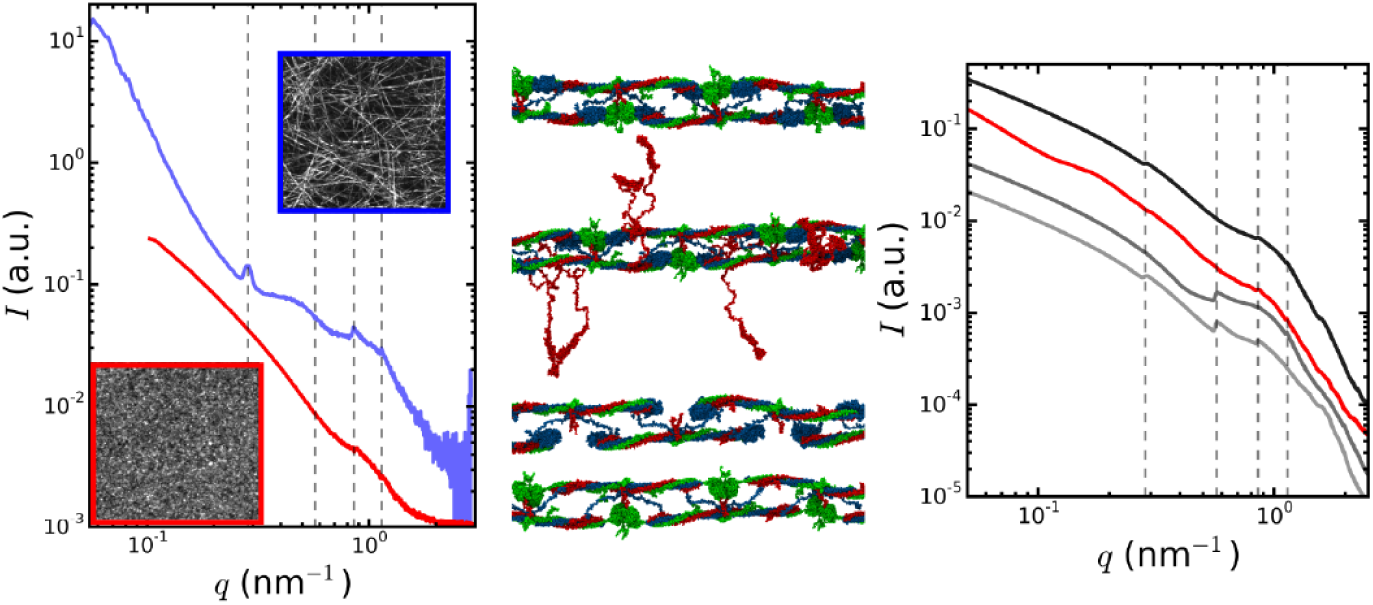

## Supplementary Information

### Supporting Table

**Table S1.** Structural characterization of the different fibrin networks used in this study. Numbers correspond to the assembly conditions as numbered in the Materials and Methods section. The fiber thickness *N*_*p*_ and protein mass density *ρ* for coarse (thick fiber) networks prepared according to conditions (2), (3) and (4) were determined by fitting the wavelength dependence of the turbidity to a theoretical model for thick fibers^1^. The data are reported as averages ± error of the mean based on at least 3 repeats per condition. For fine (thin-fiber) networks prepared according to condition (1), the wavelength-dependent turbidity was fitted to the Carr model for thin fibers^2^ to determine *ν* (average ± error of the mean based on 3 repeats). For the fiber radius, we quote an approximate range, taken from an earlier study of our lab^3^ where we imaged fine netwerks by transmission electron microscopy. The corresponding fiber density range should be regarded as indicative only given the polydispersity in *R* and the assumption of a uniform cylindrical fiber.

| Nr. | Condition details | R (nm) | $N_p$ [1] | fiber density [mg/ml], $\rho$ | mesh size ($\mu\text{m}$), $\xi$ | mass/length ($\cdot 10^{13}$ Da/cm), $\nu$ | $D_m$ |
| --- | --- | --- | --- | --- | --- | --- | --- |
| (1) | 8 mg/ml, prepared in fine clot conditions | 7.5 – 15 | $2.0 \pm 1.3$ | 71.2 – 285 | - | $0.03 \pm 0.02$ | - |
| (2) | 8 mg/ml, prepared from as-received stock | $134 \pm 87$ | $47 \pm 0.5$ | $11.6 \pm 8.9$ | $0.49 \pm 0.43$ | $0.45 \pm 0.14$ | 1.4 |
| (3a) | 4 mg/ml, prepared from dialyzed stock | $127 \pm 20$ | $291 \pm 37$ | $148 \pm 13$ | $2.66 \pm 0.30$ | $4.5 \pm 1.0$ | 1.4 |
| (3b) | 8 mg/ml, prepared from dialyzed stock | $136 \pm 5.6$ | $435 \pm 107$ | $196 \pm 60.4$ | $1.16 \pm 0.15$ | $6.6 \pm 1.6$ | 1.4 |
| (4) | 4 mg/ml, prepared from gel-filtered stock | $109 \pm 16$ | $368 \pm 52$ | $255 \pm 13$ | $1.51 \pm 0.19$ | $5.6 \pm 1.4$ | 1.5 |

## Supporting Figures

**Figure S1.**
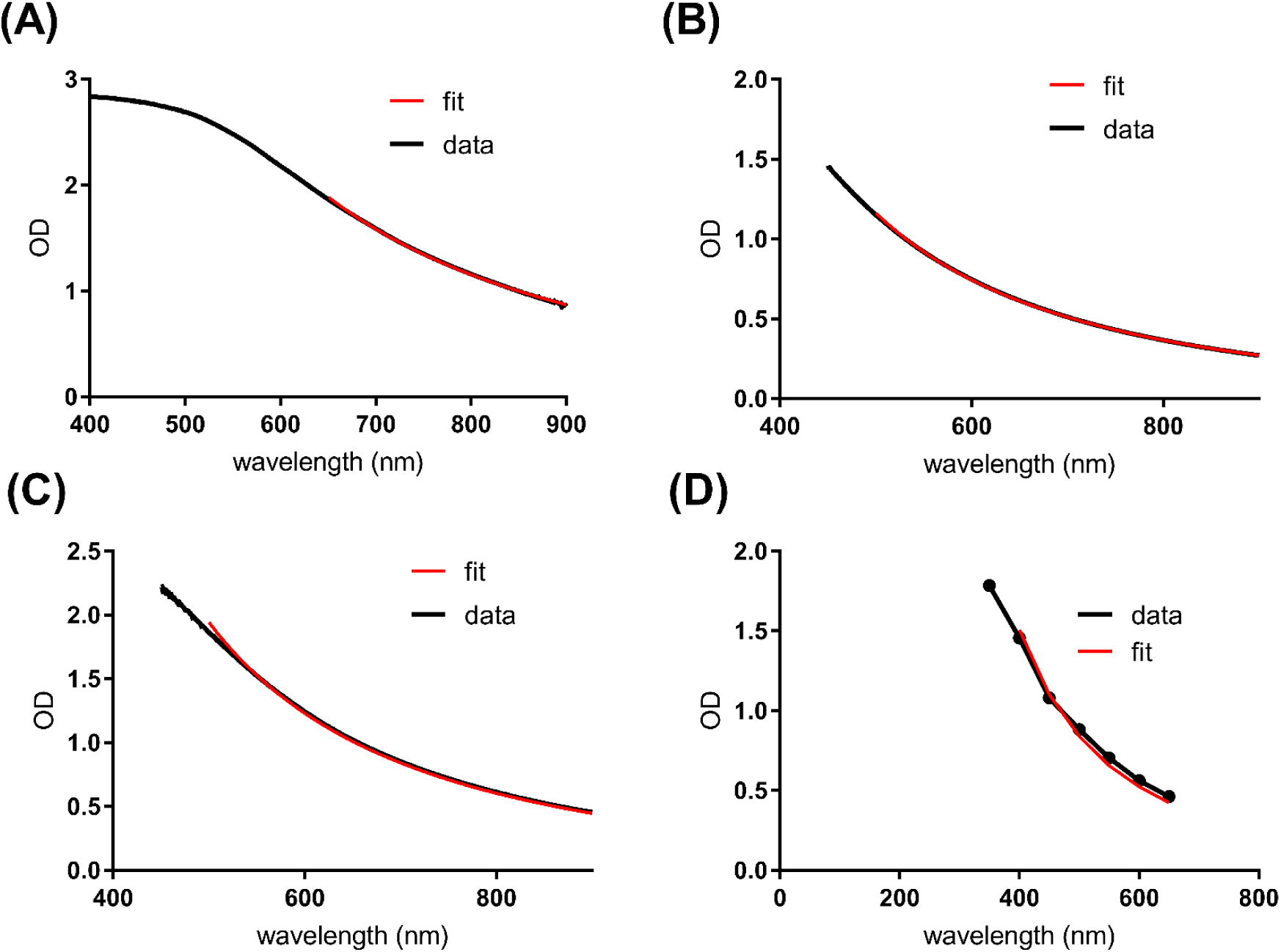
Examples of wavelength-dependent turbidity measurements (black lines) on various fibrin networks together with fits (red lines) to a theoretical model for light scattering from isotropic fibrous networks (see Materials and Methods section of the main text). The fit results are summarized in Table 1 of the main text. (A) Network prepared from dialysed fibrinogen stock (3 mg/mL). (B) Network prepared from gel-filtered fibrinogen stock (4 mg/ml). (C) Network prepared from dialysed fibrinogen stock (8 mg/ml). (D) Network prepared from as-received fibrinogen stock (8 mg/ml).

**Figure S2.**
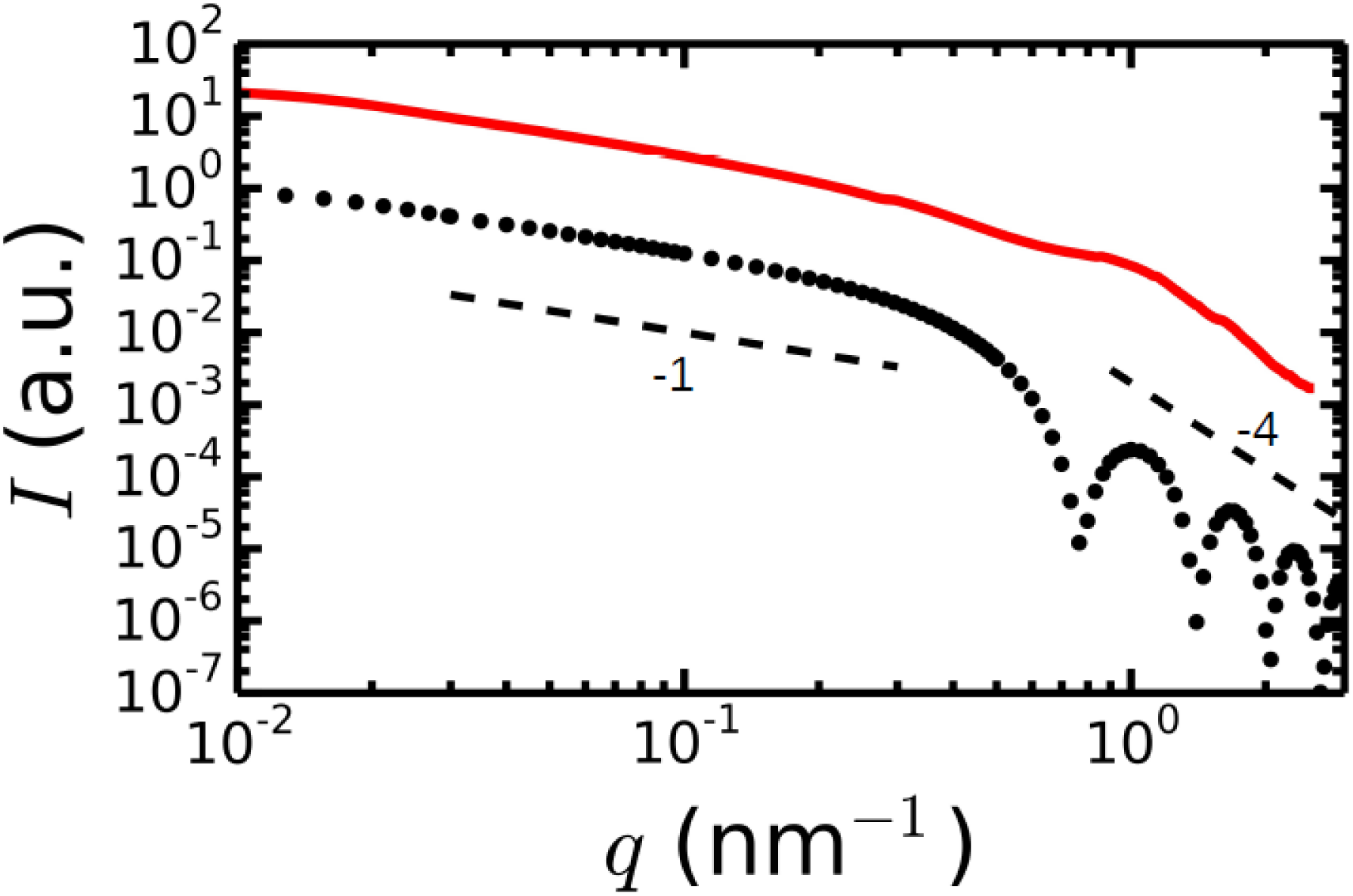
The calculated form factor of a cylinder (black circles) having the dimensions of a fibrin protofibril of 5 fibrin subunits in length (subunit length of 46 nm) and a diameter of 10 nm, compared with the predicted SAXS profile of protofibrils based on full-atom simulations of FO6-5 protofibrils (red line). Curves are shifted along the y-axis for clarity. The dashed lines indicate power laws with exponents of −1 (Guinier regime) and −4 (Porod regime), respectively.

**Figure S3.**
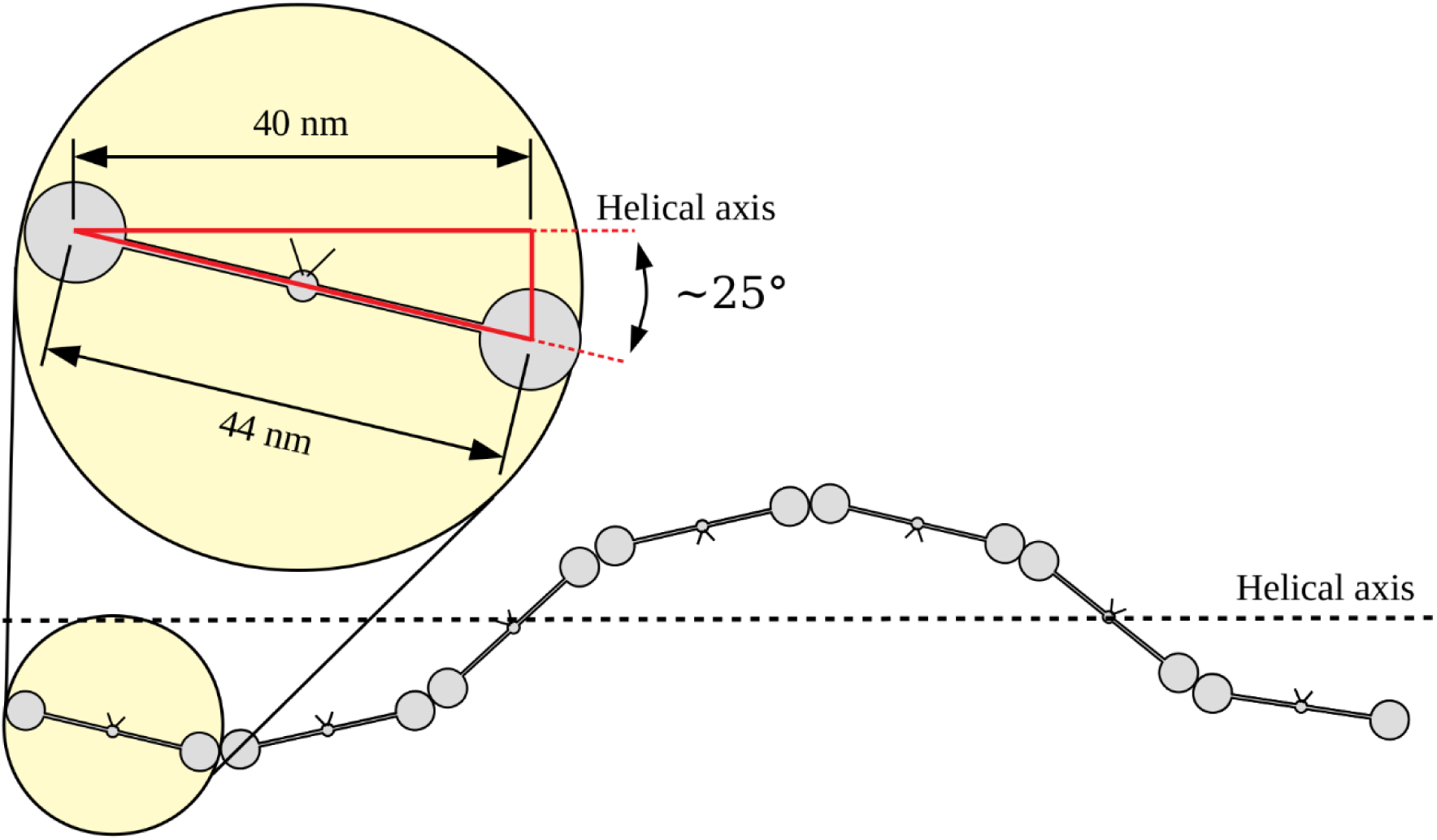
Schematic of a single stranded fibrin protofibril, highlighting the twist inside a fibrin bundle. Due to the helical structure of fibrin oligomers, the average distance between the centers-of-mass of the two globular ends of the molecule projected along the helical axis is 40 nm.

